# *De novo* motor learning of a bimanual control task over multiple days of practice

**DOI:** 10.1101/2021.10.21.465196

**Authors:** Adrian M. Haith, Christopher Yang, Jina Pakpoor, Kahori Kita

**Affiliations:** Department of Neurology, Johns Hopkins University, Baltimore, MD, USA; Department of Neuroscience, Johns Hopkins University, Baltimore, MD, USA

**Author notes:** **Corresponding Author:** Adrian Haith, 209 Carnegie, 550 North Wolfe Street, Baltimore, MD, 21287, USA.

## Abstract

Although much research on motor learning has focused on how we adapt our movements to maintain performance in the face of imposed perturbations, in many cases we must learn new skills from scratch, or *de novo*. In comparison to adaptation, relatively little is known about *de novo* learning. In part, this is because learning a new skill can involve many challenges, including learning to recognize new patterns of sensory input and generate new patterns of motor output. However, even with familiar sensory cues and well-practiced movements, the problem of quickly selecting the appropriate actions in response to the current state is challenging. Here, we devised a bimanual hand-to-cursor mapping which isolates this control problem. We find that participants initially struggled to control the cursor under this bimanual mapping, despite explicit knowledge of the mapping. Performance improved steadily over multiple days of practice, however. Participants exhibited no aftereffects when reverting to a veridical cursor, confirming that participants learned the new task *de novo*, rather than through adaptation. Corrective responses to mid-movement perturbations of the target were initially weak, but with practice, participants gradually became able to respond rapidly and robustly to these perturbations. After four days of practice, participants’ behavior under the bimanual mapping almost matched performance with a veridically mapped cursor. However, there remained a small but persistent difference in performance level. Our findings illustrate the dynamics and limitations of learning a novel controller and introduce a promising paradigm for tractably investigating this aspect of motor skill learning.

## Introduction

Much research on motor learning has focused on our ability to adapt existing skills to maintain performance in a changing environment (Shadmehr et al. 2010). This process appears to be supported by a dedicated learning mechanism, termed *adaptation*, that has been intensely studied and thoroughly characterized at the behavioral, theoretical and neural levels (Huberdeau et al. 2015b; Krakauer et al. 2019; Morehead and Xivry 2021). However, most new skills we acquire involve learning unfamiliar and often arbitrary relationships between our actions and their consequences. When driving a car, for instance, we rotate a wheel with our hands to steer and press pedals with our feet to speed up and slow down. New skills like this cannot be learned by adapting an existing controller but must instead be learned by constructing a new controller from scratch – a process which has been dubbed “*de novo*” learning (Krakauer et al. 2019; Sternad 2018; Telgen et al. 2014; Yang et al. 2021). Despite the importance of this type of learning for everyday skills, relatively little is known about it, and it remains unclear why and how controllers that are learned *de novo* might compare to pre-existing controllers.

One feature of *de novo* learning that has eluded explanation is the fact that this learning process occurs over longer timescales than adaptation. Previous studies of putative, laboratory-based *de novo* learning tasks typically require people to learn an arbitrary mapping from body position (Casadio et al. 2012; Mosier et al. 2005; Ranganathan et al. 2014), muscle activity (Berger et al. 2013; Radhakrishnan et al. 2008; de Rugy et al. 2012), or neural activity (Ganguly and Carmena 2009; Oby et al. 2019; Orsborn and Pesaran 2017; Sadtler et al. 2014) to the location of an on-screen cursor. People can learn such tasks, but learning is markedly slower than when a task can be learned through adaptation, occurring gradually over hours and days rather than in minutes. Learning of this kind results in minimal aftereffects (Ganguly and Carmena 2009; Yang et al. 2021) – in contrast to adaptation, which leads to persistent aftereffects which must be extinguished over a similar timescale to initial acquisition (Kitago et al. 2013). Finally, whereas adaptation of point-to-point movements appears to generalize strongly to online corrective movements (Ahmadi-Pajouh et al. 2012; Cluff and Scott 2013; Telgen et al. 2014), this does not seem to be the case for *de novo* learning (Gritsenko and Kalaska 2010; Kasuga et al. 2015; Telgen et al. 2014).

One potential explanation for why *de novo* learning is slow is that, because the mappings used are usually arbitrary and non-intuitive, it can be a significant challenge to discover what actions are needed to bring about a particular desired outcome. Participants may need to systematically search the space of possible actions available to them in order to identify a suitable solution, and this search process could account for why learning is so slow – particularly in high-dimensional action spaces. Another possible explanation is that it often requires participants to generate unfamiliar actions, moving fingers or activating novel combinations of muscles or neurons (Koralek et al. 2012; de Rugy et al. 2012; Sadtler et al. 2014). Learning to consistently generate these novel actions might require practice and could itself account for why overall performance improves slowly (Costa 2011; Diedrichsen and Kornysheva 2015; Oby et al. 2019).

A third possibility, however, is that even if we have explicitly identified what actions are needed to succeed at a task and can execute those actions reliably, substantial practice might still be needed to build a controller that can rapidly select an appropriate action given the current states and goals. The benefit of practice for accelerating action selection is clear in discrete-choice, arbitrary visuomotor association tasks (Hardwick et al. 2019) and assembling a controller that can select actions rapidly enough for smooth continuous control might similarly require extensive practice.

To better understand the limitations of *de novo* learning, we developed a task that isolates the problem of learning to build a novel control policy from the problems of having to discover the solution or having to reliably execute the required actions (Figure 1). The task involves maneuvering an on-screen cursor using a mapping that could easily be understood by participants (Figure 1B), thereby eliminating the problem of searching action spaces. Moreover, controlling the cursor required only planar arm movements (Figure 1A), eliminating the problem of executing unfamiliar actions. Nevertheless, participants invariably found it extremely challenging to learn to control the cursor under this mapping, requiring multiple days of practice to achieve proficient control over the cursor and to generate appropriate rapid responses to online perturbations. Crucially, the simplicity of the task also allowed us to directly compare performance to a baseline condition, providing a ceiling on the performance level that could be expected under the bimanual mapping, and revealing clear limitations on the capability of controllers learned *de novo*.

**Figure 1.**
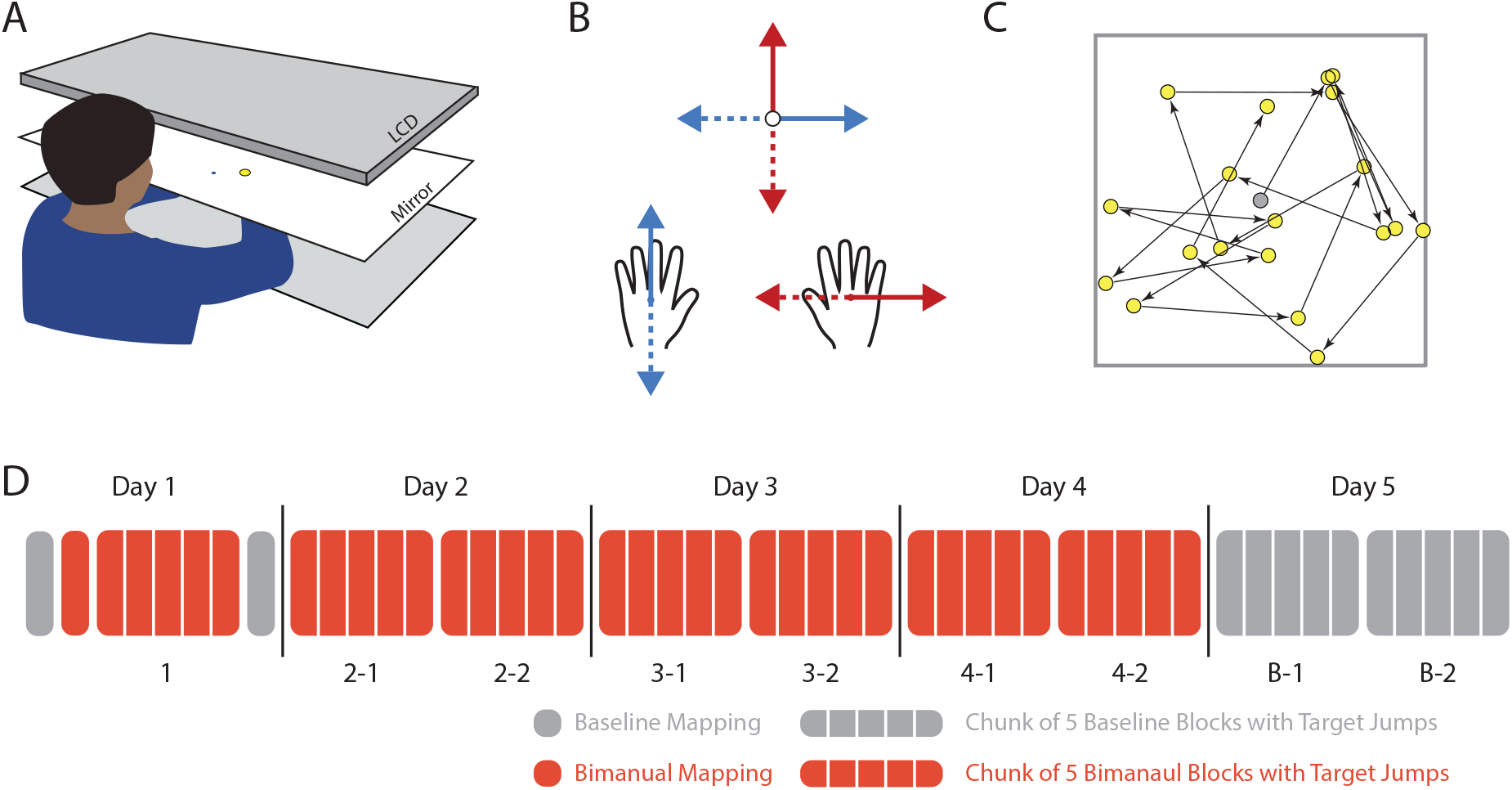
Experimental Design. A) Participants moved both of their arms on a horizontal surface to maneuver an on-screen cursor which they viewed in the plane of their arm movements via a mirrored display. B) Participants learned to maneuver a cursor via a bimanual mapping by which forward-backward movement of the left hand led to right-left movement of an onscreen cursor, while right-left movement of the right hand led to forward-backward movement of the cursor. C) In each block, participants performed a series of 60 point-to-point movements within a 20 cm × 20 cm workspace, with each new target appearing after the previous target was successfully acquired. D) Participants completed testing over 5 consecutive days, performing either Baseline blocks in which the cursor appeared at the location of the right hand (grey) or Bimanual blocks (using the Bimanual mapping in B) (red), arranged either as an isolated block with no target jumps (only on Day 1) or as a chunk of 5 blocks that included target jumps.

## Results

### Performance improves gradually over multiple days of practice

Participants learned to maneuver an on-screen cursor using a non-intuitive bimanual control interface (Figure 1B). Forward-backward movement of the left hand controlled right-left movement of the cursor, and right-left movement of the right hand controlled forward-backward movement of the cursor (the “Bimanual” mapping). This relationship between movement of the hands and movement of the cursor was explicitly described to participants. Nevertheless, they found it extremely challenging to control the cursor under this mapping. Despite the clear instructions about the task, participants often seemed to move in an exploratory manner, moving the cursor in a fixed direction until the cursor could not get closer to the target and then switching to an alternative direction (see example trajectories in Figure 2A and supplementary video) – exhibiting behavior similar to that previously described in nonlinear optimization tasks (Newell et al. 1991). During the first block of point-to-point movements using the novel Bimanual mapping, participants took, on average, over 7 seconds to complete each 12 cm movement of the cursor (Figure 2D; 7.39 s ± 1.79 s, std. dev. in mean task duration across participants), compared to less than 1 second when performing the same task using only their right hand with veridical cursor feedback (“Baseline” mapping; 0.94 s ± 0.13 s, std. dev. across participants). The trajectory of the cursor was erratic (Figure 2A), which we quantified in terms of the normalized path length (i.e., the total path length divided by distance from start to target; the lowest possible value is 1). Early trials under the Bimanual mapping had an average normalized path length of 5.99 ± 3.41 (Figure 2C; 71.8 ± 40.79 cm path length to reach a target 12 cm away), in comparison to a normalized path length of 1.20 ± 0.24 in the Baseline mapping with a veridical cursor (14.4 cm ± 2.9 cm un-normalized). Over five subsequent blocks of practice on Day 1, participants improved their performance considerably (Figure 2); by their sixth block of practice (360 trials), their average movement duration had reduced to 2.82 s ± 0.99 s, and the normalized path length reduced to near-baseline levels. Reaction times also decreased substantially.

**Figure 2.**
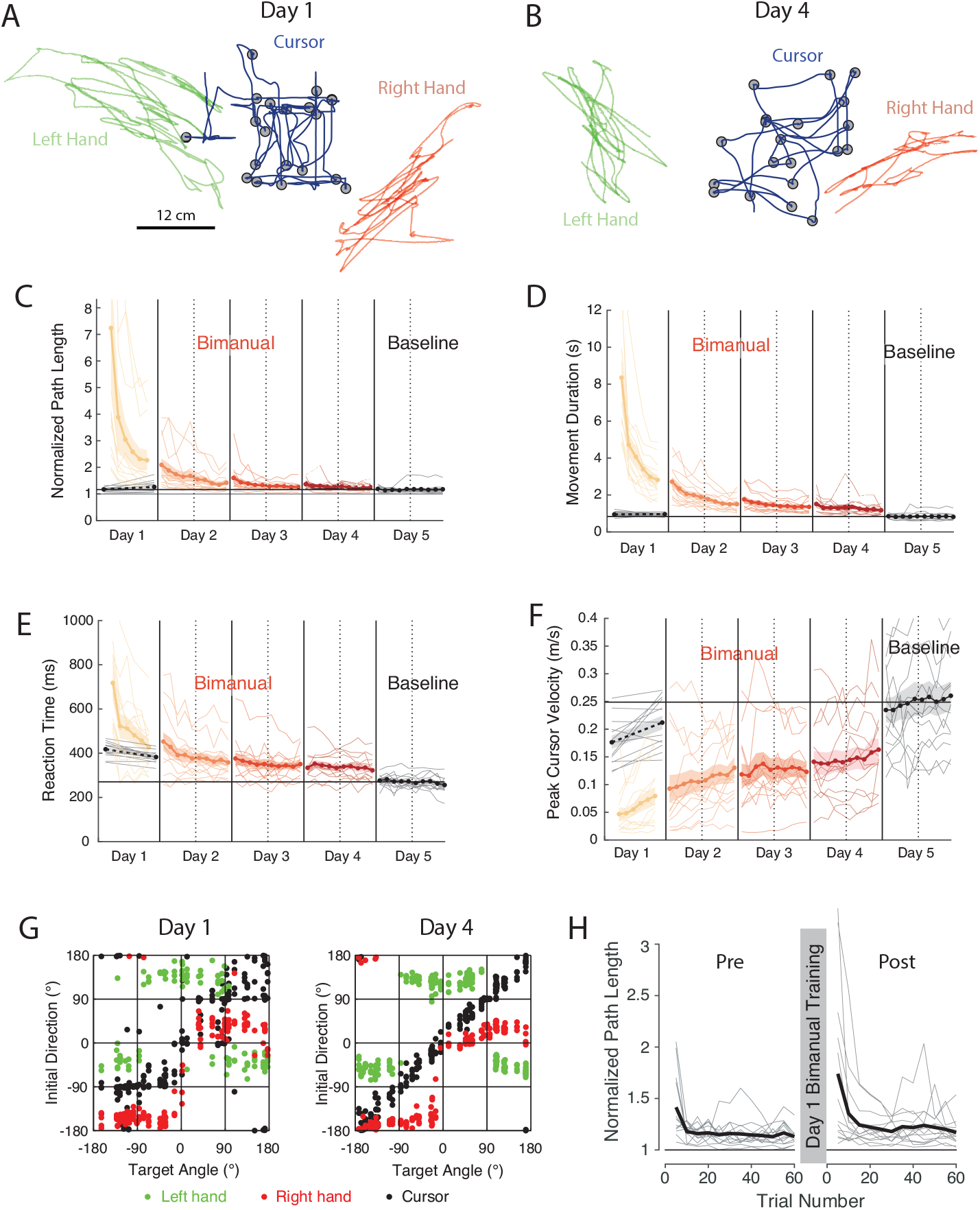
Performance under the Bimanual mapping improved gradually with practice. A) and B) Example trajectories for a representative participant in Experiment 1, showing the first 20 movements of a block during Day 1 (A) and Day 4 (B). Green and red lines show trajectories of the left and right hands, respectively (which the participant could not see), while the dark blue lines show trajectories of the cursor. C) Average normalized path length across each block for the duration of the experiment. Black points indicate performance for each block under the Baseline mapping (one before and one after practicing the Bimanual mapping). Colored lines indicate performance under the Bimanual mapping. Solid vertical lines indicate breaks between days. Vertical dashed lines indicate brief breaks between chunks of blocks. Shaded regions indicate +/- standard error in the mean across participants. D), E) and F) as C) but for movement duration, reaction time, and peak cursor velocity, respectively. G) Initial movement direction of the left hand (green dots), right hand (red dots) and cursor (black dots) in each trial on Day 1 (left) and Day 4 (right). For trials in which only one hand initially moved, only that hand is plotted. H) Normalized path length for individual trials on under the Baseline mapping on Day 1 before (left) and after (right) training under the Bimanual mapping. Thick black line indicates average across participants. Thin gray lines indicate individual participants.

To determine whether participants had learned this task through adaptation of their existing, baseline controller or by forming a brand new control policy (i.e., *de novo* learning), we removed the Bimanual mapping at the end of Day 1 and reverted back to the Baseline mapping (i.e. a cursor showing the veridical location of the right hand) to assess whether participants exhibited any aftereffects. A minority of participants initially made erroneous movements, despite having been clearly informed of the change back to a veridical cursor. However, these errors were transient rather than persistent; all participants had reverted to baseline performance levels within around 10 trials (Figure 2H). When averaged over the entire block of 60 trials, there were no significant differences in path length and movement duration between the two Baseline before and after the first 6 Bimanual practice blocks on Day 1 (all subsequent statistical hypothesis tests are two-tailed, paired t-tests unless otherwise specified; normalized path length: average difference = 0.01, p = 0.093, t(12) = 1.84, Figure 2H; duration: average difference = 14.8 ms, p = 0.73, t(12) = 0.35, Figure 2D), while reaction times *decreased* for the block after training with the Bimanual mapping than for the block before (p < .001, t(12) = 5.1, Figure 2E). Together, these results show that there were no systematic and persistent aftereffects from learning to use the Bimanual mapping, indicating that they did not learn to do so through adaptation, but rather by constructing a new controller *de novo* (Yang et al. 2021).

Participants returned to practice the Bimanual mapping for three more sessions on consecutive days, during which time their performance continued to improve steadily (Figure 2C-F). The cursor moved more directly towards the target (normalized path length: Day 2 first 5 blocks versus Day 4 last 5 blocks, p < .01, t(12) = 2.94, Figure 2C) and movement duration (p < .001, t(12) = 4.60, Figure 2D) and reaction time (p < .01, t(12) = 2.73, Figure 2E) both decreased, while movement speed (assessed based on the peak velocity over the first second of each trial) increased significantly over the same period (paired t-test: p < .01, t(12) = 3.10; Figure 2F).

By the end of Day 4, participants’ performance appeared to have reached a plateau, more closely resembling baseline levels of performance (Figure 2B, supplementary video). To test how their performance compared to baseline, we conducted a separated test session on Day 5 consisting of 10 blocks where participants controlled a veridical cursor, i.e., one that was aligned with the true location of their right hand. This test session allowed us to assess responses to target jumps under the Baseline mapping (see next section) and controlled for the possibility that participants’ performance may have improved in ways that were not directly related to learning a *de novo* controller. Performance under the Bimanual mapping at the end of Day 4 was close to performance under the Baseline mapping tested on Day 5 (Figure 2C-F). Performance did differ, however, in terms of movement duration (p < .001, t(12) = 4.6), reaction time (p < .001, t(12) = 4.70) and peak velocity (p < .01, t(12) = 3.61), but not path length (paired t-test, p = 0.19, t(12) = 1.94). Therefore, performance under the Bimanual mapping came close to that under the Baseline mapping but remained slightly poorer.

### Most participants moved their hands independently and did not rely on pre-existing coordination strategies

The bimanual mapping was designed in such a way that baseline control strategies would not have helped participants in controlling the cursor, since the required movement for each hand was orthogonal to the movements required at baseline. Nevertheless, it is possible that participants could potentially have used adapted versions of baseline control strategies to successfully control the cursor. For instance, if participants moved their hands together, the mapping would effectively have reduced to a mirror reversal of the cursor, which participants could have learned through adaptation. To characterize the degree of coordination between the hands, we examined the relationship between the initial direction of movement of the left hand and the right hand. Figure 2G shows this behavior for a representative participant both early and late in learning. For this participant, each hand primarily moved in one of two directions (“forward”/“backward” or “left”/“right”), particularly during late learning. However, these movement directions were not perfectly aligned with the direction that elicited cursor movement (+90°/-90° for the left hand, 0°,180° for the right hand). Importantly, the movement of the two hands were independent of one another, with the hands needing to move in different combinations in order to move the cursor to targets in different directions. This pattern of coordination between the hands, and its relation to the direction of the target was markedly different from baseline control in which initial hand direction is tightly yoked to target direction, as well as from any other potential coordination strategies such as yoking the two hands together or mirroring the movements of the two hands.

A minority of participants (3 of 12 in Experiment 1 and 1 of 8 in Experiment 2), however, did seem to adopt a different strategy in which their hand direction for one hand was more continuously tuned to the target direction, aimed approximately 90° counterclockwise from the true target (See participants 1-3, 1-5, and 1-9 in Figure S1, and 2-4 in Figure S2). This pattern could potentially have been attributable to adaptation of a baseline controller, for one hand at least. Nevertheless, although there was some heterogeneity in the way participants solved the task, a large majority of participants solved the task in a similar manner, moving their two hands independently, not relying on any pre-existing coordination between the hands or adaptation of pre-existing control strategies.

### Rapid feedback responses emerged gradually with practice

In addition to quantifying the quality of participants’ point-to-point movements, we also examined participants’ ability to generate rapid corrections when the target jumped to a new location during movement. In 1/3 of trials, the target was displaced orthogonal to the required movement direction by either +/- 1.5 cm or +/- 3 cm, after the cursor crossed a line 1/3 of the distance between the start position and the target (Figure 3A). On the first day of practice, participants generated weak and erratic responses to target jumps (Figure 3B,C). Starting on the second day, however, a clear response emerged which became more rapid over subsequent days of practice (Figure 3B,C).

**Figure 3.**
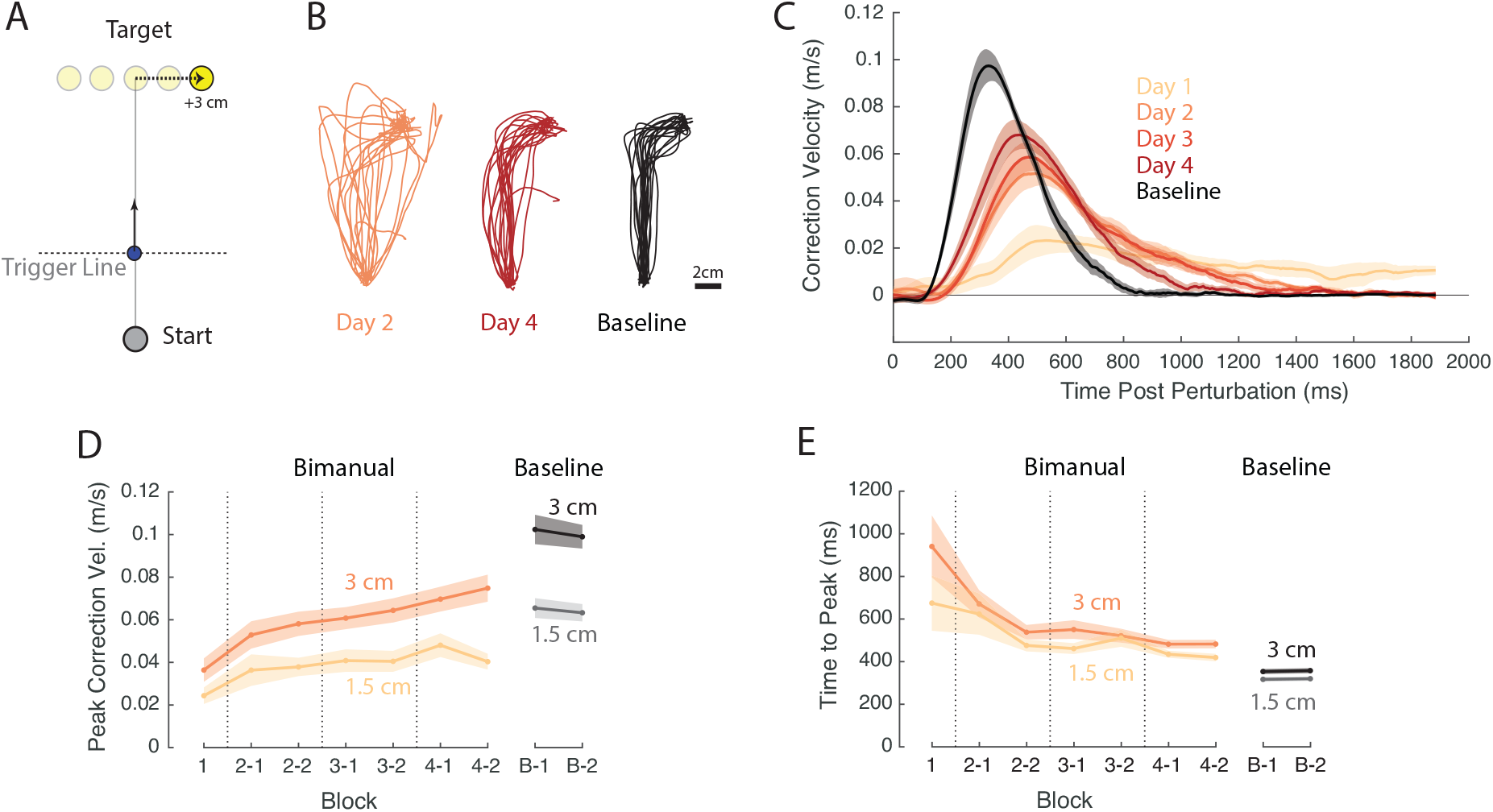
Emergence of rapid feedback responses with practice. A) Within certain blocks, a subset (1/3) of targets could jump orthogonal to the primary direction of movement by 1.5 cm or 3 cm during movement, prompting a corrective response. Jumping of the target was triggered when the cursor crossed an invisible line 1/3 of the way to the original target location from the start position. B) Example trajectories from a representative participant following a 3cm rightward target jump early in learning (Day 2; orange), late in learning (Day 4, red) and under the baseline mapping (black). Behavior from Day 2 plotted here since Day 1 behavior was very erratic for all participants. C) Velocity profile of corrective responses in the direction parallel to the target jump for 3cm jumps, averaged across jump directions (flipped accordingly). Colored lines indicate corrective response under the Bimanual Mapping at different stages of learning based on performance from the last 5-block chunk on each day (i.e. chunks 1, 2-2, 3-2, 4-2). Black line indicates corrective response under the Baseline Mapping. D) Peak correction velocity as a function of practice under the Bimanual mapping (colored lines) and under the Baseline mapping (black/gray lines). Error bars show +/- standard error in the mean across participants. E) As D) but showing the latency to peak correction velocity.

We quantified the strength of the feedback response in terms of the peak correction speed (parallel to the target displacement) and the latency to this peak speed. The peak speed of correction increased from the first to the fourth day for both large and small target jumps (Figure 3D; 1.5 cm jump: p < .005, t(12) = 4.15; 3cm jump: p < .0001, t(12) = 6.39) and the latency to this peak also decreased (Figure 3E; 1.5 cm jump: p < .0001, t(12) = 6.39; 3 cm jump: p = 0.014, t(12) = 4.15). However, the velocity of corrective movements remained slower than those under the Baseline mapping (1.5 cm jump: p < .005, t(12) = 3.94; 3 cm jump: p < .0001, t(12) = 6.39), and the peak velocity occurred later (1.5 cm jump: p < .0001, t(12) = 6.39; 3 cm jump: p = 0.002, t(12) = 3.93) suggesting that, even though the *de novo* skill had been well learned, performance was still substantially worse than their baseline performance. We also examined whether the latency of response onset differed across mappings or with extent of practice. We estimated response onset by fitting a “template” response (a piecewise linear-quadratic function) to the early part of the response velocity profile (see Materials and Methods for details). We used this approach rather than alternative approaches based on signal-detection since, under these approaches, the estimated time of response onset can be strongly biased by the strength or weakness of the response, which we expected to differ significantly across conditions. We found that, under the Baseline mapping, online corrective responses began at 186 ms and 196 ms after the target jump for small and large target jumps, respectively (average of both blocks on Day 5). Under the Bimanual mapping, by contrast, responses began at 223 ms and 233 ms for small and large target jumps, significantly slower than under the Baseline mapping (1.5 cm jump: p = 0.032, t(12) = 2.417; 3 cm jump: p = 0.043, t = 2.253).

### Slower feedback corrections under the Bimanual mapping were not attributable to slower primary movements

A limitation of Experiment 1 is that participants typically moved more slowly under the Bimanual mapping than under the Baseline mapping (t-test on peak velocity of primary movement: p = 0.0034, t(7) = 3.64). This difference in the speed of primary movements might have accounted for the slower feedback corrections under the Bimanual mapping, particularly given that the speed of movement preparation has been linked to movement vigor (Kita et al. 2022; Thura et al. 2014). We conducted a second experiment in which, starting on Day 2, we gave participants feedback about the peak velocity of the cursor at the end of each movement (see Methods), enabling them to achieve more consistent movement speeds across the Baseline and Bimanual mappings (Figure 4A,B; peak velocity on last day of Bimanual practice vs Baseline: p = 0.821, t(7) = 0.235). As in Experiment 1, participants improved their performance, assessed in terms of path length, over multiple days of practice (Figure 4C). Note that the deterioration in performance between the end of Day 1 and the start of Day 2 was due to speed requirements being introduced at the start of Day 2. Despite the speed of the primary movement being matched across mappings, the speed of feedback corrections remained significantly slower under the Bimanual mapping than under the Baseline mapping, even after 3 days of practice (Figure 4E; 1.5 cm jump: p < 10^−5^, t(7) = 11.49; 3 cm jump: p < .01, t(7) = 4.35). The latency of peak responses was also greater for the Bimanual mapping compared to under the Baseline mapping (Figure 4F; 1.5 cm jump: p< 10^−7^, t(7) = 25.4; 3 cm jump: p < 10^−7^, t(7) = 23.7), as was the latency of response onset (1.5 cm jump: p = 0.009, t = 3.58; 3 cm jump: p = 0.0014, t = 5.12; not shown). Therefore, even when the vigor of primary movements was well matched across conditions, participants could not correct errors as rapidly under the *de novo* controller learned for the Bimanual mapping.

**Figure 4.**
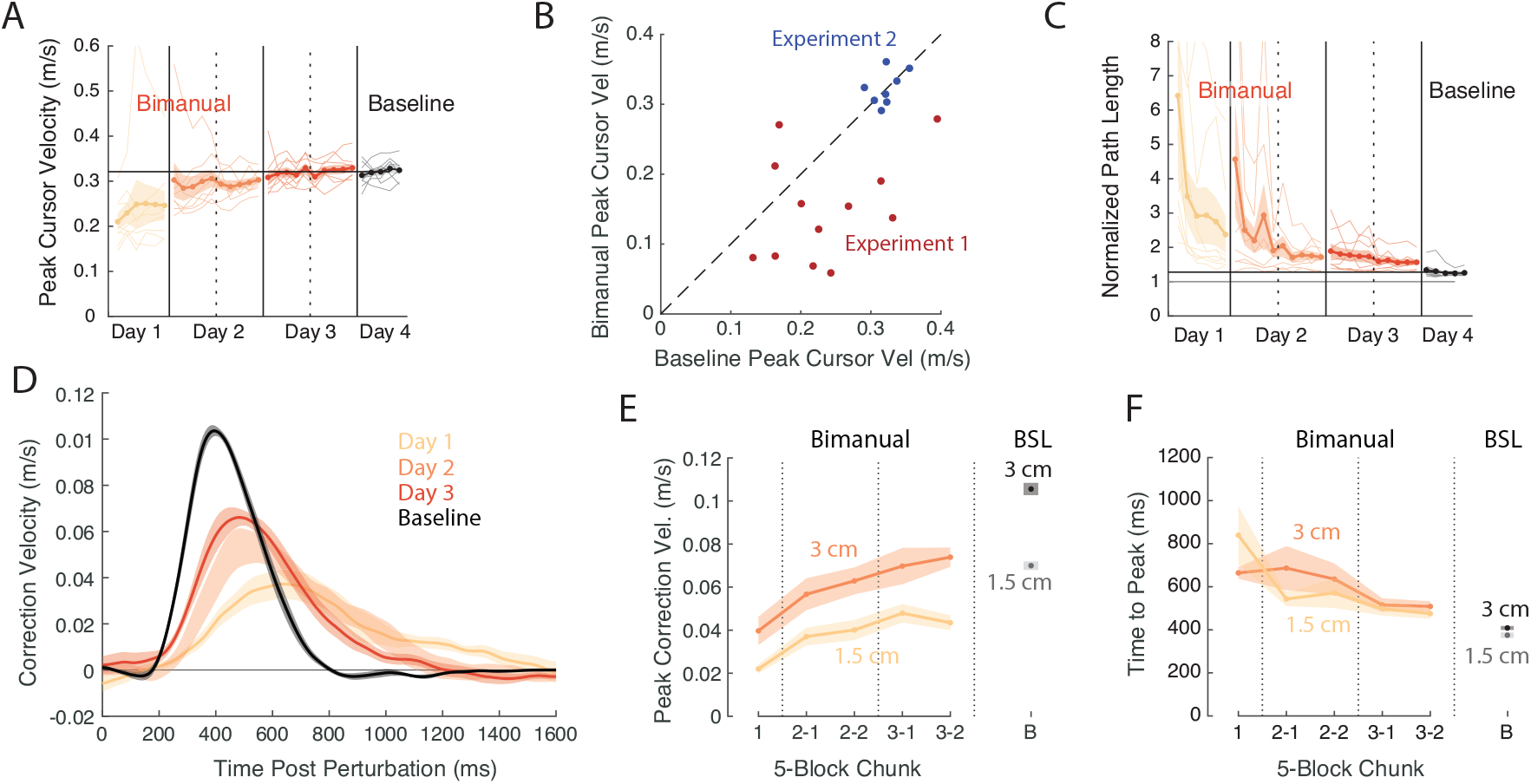
Results from Experiment 2 in which movement speed was controlled by providing participants with feedback about the peak cursor velocity. A) Average peak cursor velocity across each block for the duration of Experiment 2 (as Figure 2F). Peak cursor velocity was well-matched between the Bimanual and Baseline conditions. B) Comparison of peak velocity under the Baseline mapping and in the final block of practice under the Bimanual mapping for Experiment 1 (red dots) and Experiment 2 (blue dots). Each dot represents an individual participant. C) as A), except for normalized path length. D) Averaged velocity profiles of corrective movements following target jumps in Experiment 2 (as Figure 3C). E) Peak correction velocity (as Figure 3D). F) Latency to peak correction velocity (as Figure 3E).

### Slow feedback corrections under the Bimanual mapping were not due to controlling two hands rather than one

One possible explanation for why participants were able to respond more rapidly under the Baseline mapping than under the Bimanual mapping is that the Bimanual mapping required them to use both hands, whereas the Baseline mapping only required them to use their dominant right hand. The slower response under the Bimanual mapping could conceivably be attributable to having to coordinate the use of both effectors, or might reflect slower responses of the non-dominant hand. To test whether the use of both hands versus a single hand could account for the discrepancy in behavior, we performed an additional experiment (Experiment 3) in which participants controlled the cursor either with their dominant (right) hand, non-dominant (left) hand, or both hands together with the cursor appearing at the average position of the two hands. Participants’ corrections to target jumps did appear slightly faster when using their right hand alone in comparison to using both hands to control the cursor (Figure 5A). The peak velocity did not differ significantly according to how the cursor was controlled (ANOVA, p = 0.07, F(2,54) = 2.77), but the latency to peak correction did (ANOVA, p = 0.016, F(2,54) = 4.47). Critically, however, the differences in correction velocity when using the right hand versus using both hands in Experiment 3, where the mapping was veridical in both cases, was significantly less than the same difference for Experiment 2 (two-way ANOVA, main effect of experiment: F(1,32) = 28.94, p < .0001), and the same was true for the latency to peak velocity (two-way ANOVA, main effect of experiment: F(1,32) = 16.76, p = 0.0003). These data demonstrate that even if the use of both hands causes movements to slow down compared to movements with just the dominant hand, this difference cannot entirely account for the difference in performance between the Baseline and Bimanual mappings in Experiment 2. Instead, the slower feedback corrections under the Bimanual mapping were a property of the learned controller.

**Figure 5.**
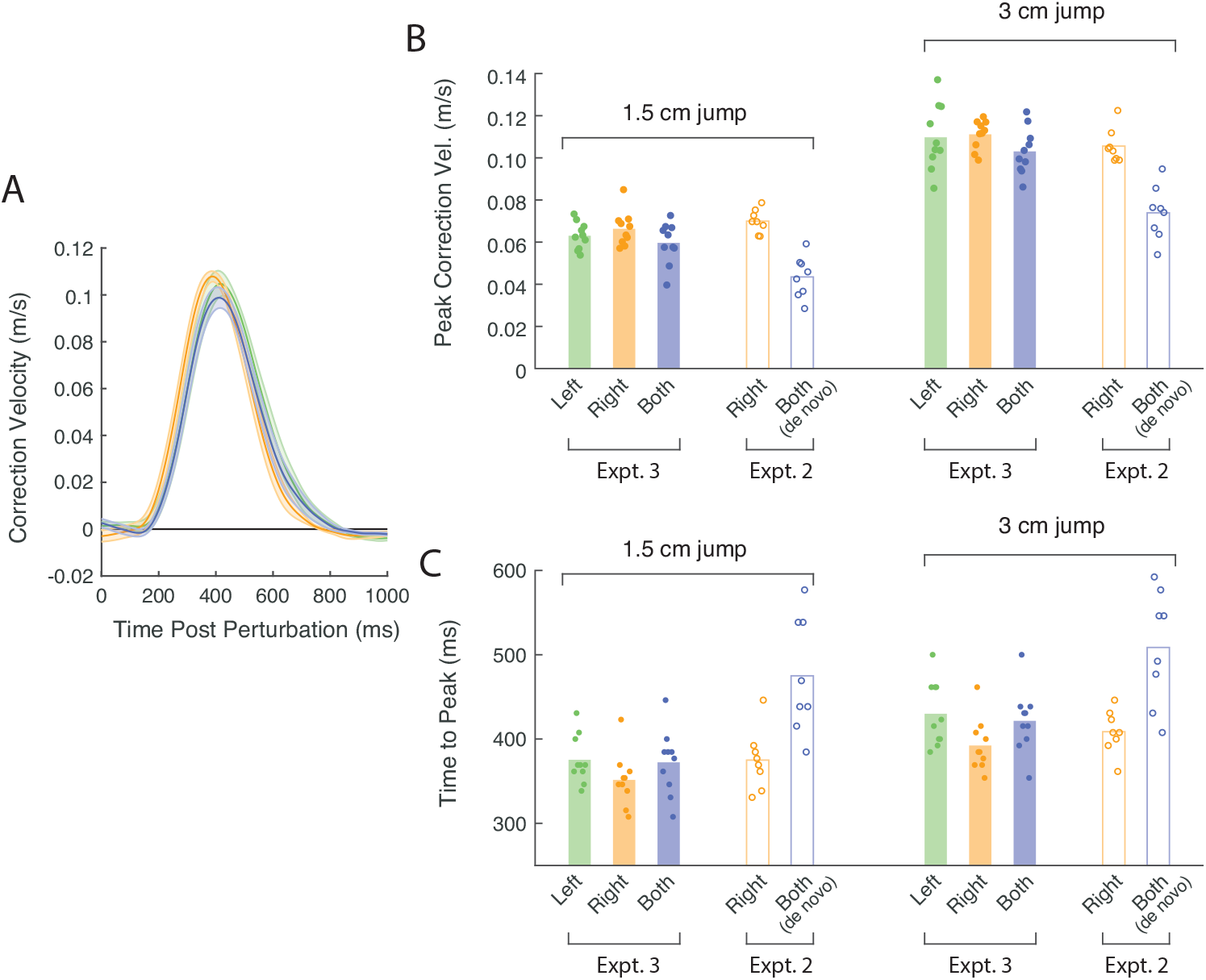
Using one hand versus two hands to maneuver the cursor does not account for slow response speed under the Bimanual mapping. A) Velocity profile of corrective responses parallel to the direction of the target jump in the left-hand (green), right-hand (orange), and both-hands (blue) conditions, as in Figure 3B. B)-C). Peak correction velocity and latency to peak correction velocity for different handedness conditions and jump sizes in Experiment 3 (filled markers and bars) and Experiment 2 (unfilled markers and bars). Dots represent behavior of individual participants. Colored bars indicate mean behavior across participants.

## Discussion

We have introduced a novel motor learning paradigm for studying *de novo* motor learning. Our paradigm is simple in that it involves only a linear mapping from planar arm movements to cursor movements that can be easily explained to participants. However, the mapping itself is very challenging to learn, requiring multiple sessions of practice to gain proficiency. The use of planar arm movements means that participants do need to learn to generate unfamiliar movements, as is often the case in many kinematic, myo-electric and BMI-based learning tasks. Furthermore, by testing participants’ behavior in the same tasks but with a veridical cursor (the Baseline mapping), we are able to establish a ceiling on the performance level that could be expected of participants in this task. Our results clearly demonstrate gradual improvements in performance over multiple sessions of practice, and also show that, even after 4 days of training, participants’ performance under the Bimanual mapping still remained worse than under the Baseline mapping.

In principle, learning to control a cursor under our Bimanual mapping could be construed as countering a perturbation, rather than as acquiring a new skill. However, the absence of persistent aftereffects demonstrates that participants did not learn the task through adaptation mechanisms, but instead built a new policy, i.e. through “*de novo*” learning (Yang et al. 2021). Although some aftereffects seemed to be present when participants switched from the bimanual mapping back to the baseline mapping, these aftereffects were only apparent in a subset of participants. Furthermore, these aftereffects lasted for only around 10 trials, despite these trials occurring to targets in many different directions. In contrast, aftereffects from visuomotor rotation adaptation can persist for up to 30 trials to a single target (Kitago et al. 2013). The lack of persistent aftereffects suggests that the learning we observed here might be similar to that seen in response to mirror reversals of visual feedback (Lillicrap et al. 2013; Telgen et al. 2014; Yang et al. 2021). However, an important difference between our Bimanual mapping and a mirror reversal is that, under a mirror reversal, the required movement under the reversal is in direct conflict with the movement that would be required at baseline. By contrast, under the Bimanual mapping, the hand movements required under the Bimanual mapping are orthogonal to those required at baseline. Moreover, a mirror-reversal can be solved via a simple re-aiming strategy (Wilterson and Taylor 2021), whereas such a strategy is not readily available under the Bimanual mapping. Closely examining the way in which participants coordinated their two hands to move the cursor in different directions (Figure 2G, S1 and S2) confirmed that, for the majority of participants, they learned to maneuver the cursor by coordinating their two hands in a very different way from baseline patterns of control. A small minority of participants, however, did appear to adopt a different solution in which movement of one hand was rotated 90° relative to the target direction (rather than strictly just left/right), suggesting that they might have leveraged pre-existing controllers to some extent. Nevertheless, the majority of participants did not appear to employ such an approach.

We argued that participants learned to control the cursor via *de novo* learning and not via adaptation based on the lack of aftereffects when switching between them in Experiment 1. A potential concern with this logic is that the lack of aftereffects might be attributable to a contextual switch from using two hands to one hand. However, numerous studies have found that adaptation generalizes partially but robustly from unimanual to bimanual movements (Burgess et al. 2007; Nozaki et al. 2006; Wang et al. 2013; Wang and Sainburg 2009). Therefore, we think it very unlikely that participants could have learned the Bimanual mapping using adaptation mechanisms but failed to exhibit aftereffects due to the switch to unimanual movements. Nevertheless, we suggest that for future investigations, the bimanual-average mapping in which the cursor appeared at the average location of the two hands, as we used in Experiment 3, would serve as a more suitable baseline mapping for comparison to behavior under the Bimanual mapping.

A major question not resolved by our present experiments is whether, with additional practice, performance under the Bimanual mapping could eventually match that under the Baseline mapping, both in terms of the quality of point-to-point movements and of rapid corrective responses. In a small set of participants tested for longer periods, we have found that 10 days of practice led to minimal further improvements (Yang et al. 2022). Anecdotally, we found that two authors of this study who practiced the Bimanual mapping extensively (∼3 weeks) did not appear to exhibit any performance gains beyond those seen in the results of Experiment 1. A larger cohort will be required to rigorously test whether this is the case in general. It is possible that extremely rapid responses using a veridical cursor is achieved by engaging rapid subcortical pathways (Perfiliev et al. 2010), whereas using the Bimanual mapping requires engagement of slower cortical circuitry, accounting for the seeming limit on performance under the Bimanual mapping. However, the extent of subcortical involvement in simple arm movements in humans remains unclear.

Another important question not answered by our current experiments is *how* participants learned to control the cursor under the Bimanual mapping. Our results do show clearly that explicit knowledge of the mapping is insufficient to perform the task (since the mapping was explained to all participants as it was first introduced), highlighting a critical role of practice in learning to skillfully control the cursor. It is well established that cognitive processes play a critical role during adaptation (McDougle et al. 2016; Seidler et al. 2012), and we expect that this is equally true, if not more so, in the case of *de novo* learning. Consequently, many themes related to cognitive aspects of adaptation might also prove to be germane in the case of *de novo* learning, including deliberative strategies (McDougle and Taylor 2018; Morehead et al. 2015), retrieval (Huberdeau et al. 2015a; Kumar et al. 2021), and temporal stability of memories (Bindra et al. 2021; Hadjiosif and Smith 2013; Sing et al. 2009). Other, less cognitive, potential learning mechanisms for *de novo* learning have also been proposed, based on forward and inverse models (Pierella et al. 2019) or based on optimization in artificial neural networks (Hennig et al. 2021). We have recently found that, in this same Bimanual task, performance becomes habitual early within 1-2 days of practice (Yang et al. 2022), suggesting that the bulk of improvement over days might occur through implicit, rather explicit learning mechanisms. We suggest that identifying and validating potential learning algorithms for this type of task ought to be a major focus of future motor learning research.

A number of paradigms have been devised to study motor learning that appear to engage the same or similar kind of learning as we consider here. These include body-machine interfaces using either movements of the hand (Mosier et al. 2005; Ranganathan et al. 2014) or other parts of the body (Casadio et al. 2012) to maneuver a cursor. Work on brain-machine interfaces (BMIs) also challenges humans or monkeys to learn to maneuver an on-screen cursor and, despite best efforts to calibrate the decoder to match the subject’s natural movement, some degree of learning is also known to occur (Carmena et al. 2003). BMI-based paradigms have, consequently, been leveraged to achieve basic insights into motor learning, often by imposing arbitrary relationships between neural activity and movement of a cursor (Golub et al. 2016; Koralek et al. 2012; Orsborn and Carmena 2013; Sadtler et al. 2014). In more abstract tasks, participants do not directly control a cursor but are simply required to minimize an unknown function of their hand positions (Krinskii and Shik 1964; Newell et al. 1991). Such tasks examine how participants converge on the solution through exploration and, although they don’t examine long-term practice-related improvement that is present in other *de novo* learning paradigms, we believe that the nature of behavior in such tasks is likely very similar to participants’ performance in the very early stages of learning our bimanual cursor control task.

The specific paradigm presented here was devised to isolate the problem of learning a new mapping from familiar sensory states and cues (the location of an on-screen cursor and target) to well-practiced motor output (planar arm movement). More broadly, however, *de novo* learning might encompass learning how to generate commands based on novel sensory input (Bach-y-Rita and W. Kercel 2003; van Vugt and Ostry 2017) or generating patterns of motor output that one has never had to generate before (Costa 2011; Oby et al. 2019; Sadtler et al. 2014). Learning most new skills likely involves a combination of all three challenges. In particular, studies of motor learning that challenge participants to learn to control a novel brain-machine interface often requires them to generate novel patterns of motor output (Sadtler et al. 2014), in which case learning is significantly slower and engages alternative neural mechanisms (Oby et al. 2019). The relative simplicity of our task allowed participants to exploit existing patterns of coordination (for instance, moving the arms in tandem (Kelso et al. 1979)) to achieve proficient performance. Yet, despite the fact that there was considerably less to learn than in other *de novo* skill learning tasks, participants still found the task extremely difficult and learned slowly. Our findings here demonstrate that learning to select specific actions in the right context also presents a challenging problem and research using this and similar paradigms can provide a complementary approach to BMI-based research on learning.

In recent years, much research in the computational motor learning community – particularly behavioral studies in human subjects – has been dominated by the study of adaptation. In adaptation tasks, an already well-learned behavior (such as reaching, or saccades) is systematically perturbed and participants must learn to counter the perturbation in order to regain prior levels of performance. Adaptation is, however, unlikely to provide a good model of how we acquire new skills. Adaptation usually occurs in minutes, whereas learning a new skill such as riding a bicycle, juggling, or playing an instrument can take days, weeks or months. We believe that the kind of “*de novo*” learning studied here provides a more suitable model of at least certain aspects of skill acquisition, compared to adaptation and ought to become a more major focus of motor learning research.

We suspect there are a number of reasons why *de novo* learning has been relatively understudied in comparison to adaptation. First, adaptation is rapid, often occurring in just minutes. This makes it pragmatically very accessible to study in comparison to learning tasks in which learning might need to occur over multiple sessions. Second, behavior in adaptation paradigms can be easily quantified in terms of the initial direction of a reaching movement or of an applied force. These metrics also seem to work well as the putative basis of learning – computational models of adaptation can predict trial-by-trial changes in reach direction remarkably accurately. It therefore seems reasonable that, in a task like visuomotor rotation, the motor system does indeed “learn” the appropriate reach direction. By contrast, in a *de novo* learning task, participants do not “learn” an appropriate path length or movement duration. These gross metrics are far-removed from the underlying representational changes in the brain. Rather, they are emergent properties of a learned policy. In adaptation tasks, the space of learned policies (and, thus, the state of learning) appears to be well parametrized by an angle parameter – or by a series of such parameters learned by different processes (Huberdeau et al. 2015b; Kording et al. 2007; Morehead and Xivry 2021). In the case of *de novo* learning, it is unclear how the space of potential policies ought to be parametrized in order to appropriately characterize both the state of learning at any given time and the dynamics by which learning proceeds from trial to trial.

In our experiments, we attempted to gain more insight into the underlying controller being learned by probing rapid motor responses to correct for mid-movement target jumps. These target jumps revealed gradual improvements in the speed of corrective responses with practice. From an engineering perspective, target jumps provide a means of probing the control system that, in principle, allow us to comprehensively characterize its properties. For a linear control system, knowing how the system responds to target jumps should be sufficient to predict the response to arbitrary movements of the target. Occasional target jumps are an inefficient experimental approach, however. There do exist alternative, more powerful, approaches to characterizing controllers based on system identification in tracking tasks. These methods are considerably more data-efficient than the target-jump approach, and have been successfully applied to understand basic properties of control (Yamagami et al. 2019; Zimmet et al. 2020) and learning of visuomotor gains, rotations and mirror reversals (Yang et al. 2021; Zimmet et al. 2020). In future work, we plan to apply this approach to try to understand learning in more complex *de novo* learning tasks like the one presented here.

## Methods

A total of 31 participants (9 male, 22 female; average age 24.7) were recruited for this study. Participants provided written consent, and all study procedures were approved by the Johns Hopkins University School of Medicine Institutional Review Board.

### Experiment 1

We recruited 13 participants for Experiment 1 (3 male, 10 female; average age 24.2). Participants sat at a glass-surfaced table, with both arms supported on pressurized air sleds that enabled frictionless planar movements of the arms. Participants could not directly see their arms, but viewed a horizontally mounted, downward-facing monitor through a horizontal mirror, positioned such that the image of the monitor appeared in the plane of movement (Figure 1A). Hand position was tracked at 130 Hz using a Flock of Birds tracking system (Ascension Technology).

### Point-to-point task

In each block, participants moved their hands to maneuver an on-screen cursor (5 mm diameter) to a series of targets. To start the block, participants were instructed to move the cursor to a circle (10 mm diameter) appearing in the center of the workspace. Participants were given no specific instructions about how quickly they should move the cursor in Experiment 1. Once the participant held the cursor stationary inside this circle, it moved to a new location 12 cm away. After participants acquired this target, it moved again. The distance between successive targets was always 12 cm, but the direction was selected pseudorandomly, subject to the constraint that all targets had to fall within a 20 cm × 20 cm workspace centered on the start position. A single block consisted of 60 targets (not counting the initial starting circle). Participants performed this task either under the Baseline mapping, in which the cursor appeared at the true position of their right hand, or under the Bimanual mapping. Under the Bimanual mapping, the x-location of the cursor was determined by the y-location of their left hand and the y-location of the cursor was determined by the x-location of their right hand. Changes in the x-location of the left hand and changes in the y-location of the right hand had no effect on the location of the cursor. This mapping was explicitly explained to participants prior to the start of their first time experiencing the Bimanual mapping.

### Target Jumps

In some blocks, we probed online rapid responses with target jumps. One third of targets (non-consecutive) in a given block were selected as targets that could jump. During movements to these targets, the target would jump once the cursor crossed a perpendicular line one third of the distance to the target (Figure 3A) by ± 1.5 cm, ± 3 cm, or 0 cm. Participants completed blocks in chunks of 5 with the same target sequence each time, experiencing each jump amplitude once at each jump target, in a different pseudorandom order across trials for each target. Thus, within each chunk of 5 blocks, participants experienced 20 jump trials of each possible size.

### Experiment Structure

Participants first completed an initial block of 60 movements under the Baseline mapping without target jumps. They then completed a block of 60 movemetns under the Bimanual mapping without target jumps, after which they completed a chunk of five 60-trial blocks with target jumps, structured as described above. For the final block on the first day, participants performed a single block of movements using just their right hand, with veridical feedback, to assess any potential aftereffects from learning to control the cursor under the Bimanual mapping. Participants returned to practice the Bimanual for three further sessions on consecutive days. In each of these sessions, they completed two chunks of five blocks (600 trials total) that included target jumps, which lasted approximately half an hour. On Day 5, participants returned to complete two more chunks of five blocks (600 trials total), this time using the Baseline mapping in which the cursor was aligned with the true position of the right hand.

### Experiment 2

In Experiment 2, we sought to better control the speed of participants movements across the Baseline and Bimanual mappings. We recruited 8 participants for this experiment (3 male, 5 female; average age 23.8). Participants were instructed to try to keep the speed of their movements consistent throughout the experiment. After each target was acquired, we provided participants feedback about the peak velocity of their movement. If the peak speed of the cursor was less than 0.3 m/s, the target they had just reached turned blue, indicating that the movement was too slow. If the peak speed of the cursor was greater than 0.4 m/s, the target turned red, indicating that the movement was too fast. If the peak speed was between 0.3 m/s and 0.4 m/s, the target turned green, indicating a good speed.

Experiment 2 followed a similar structure to Experiment 1, except participants only practiced the Bimanual mapping for 3 days instead of 4. Feedback about movement speed was not given on Day 1, since it was very difficult to comply with this requirement while initially learning the Bimanual mapping. Feedback about movement speed was provided on Day 2 onwards.

### Experiment 3

In order to determine whether the differences we observed in feedback corrections between the Baseline and Bimanual mappings was due to using one hand or two hands, we performed an additional control experiment to assess performance using different effectors. Ten participants (3 male, 7 female; average age 26.1) completed 3 chunks of 5 blocks with target jumps and feedback about movement speed. For each chunk, the cursor was aligned either to: i) the veridical location of the participant’s left hand, ii) the veridical location of the participant’s right hand, or iii) the location halfway between their left hand and right hand.

### Data analysis

Position data of both hands and the cursor were smoothed using a Savitzky-Golay filter to eliminate high-frequency noise (3^rd^-order; window size = 7 samples / 5.3 ms) and numerically differentiated to obtain movement velocity. These velocity signals were then smoothed again using the same approach. The movement onset time was determined based on the first timepoint when the cursor’s movement velocity exceeded 0.025 m/s after the target was presented and this was also used to determine the reaction time. The movement end time was determined based on the earliest time at which participants successfully held the center of the cursor stationary (tangential velocity < 0.025 m/s) within the target. Movement duration was determined based on the difference between movement onset and movement end time. The path length was computed as the total length of the smoothed trajectory between the movement onset time and the movement end time. The normalized path length was this distance divided by the distance between the center of the start position and the target position (12 cm). Only movements in which the target did not jump were included in the path length analysis. Differences in performance between mappings and between different points during learning were assessed using paired t-tests.

To analyze corrective movements in response to target jumps, we extracted the velocity of the cursor parallel to the direction of the target jump (perpendicular to the straight line between the start position and initial target position for each movement) and aligned these trajectories to the time at which the target jumped. We averaged velocity profiles for leftward and rightward jumps of the same magnitude by flipping the sign of the data for responses to leftward target jumps. To quantify corrections, we computed the peak correction velocity for each trial and averaged these across jumps of similar size within each chunk of 5 blocks. We also computed the average time at which this peak correction velocity was attained.

We estimated the time at which the response to the target jump was first initiated by fitting the averaged velocity profiles using a simple model of the corrective velocity profile consisting of a linear portion before the onset of the response, and a quadratic portion after the response was initiated:

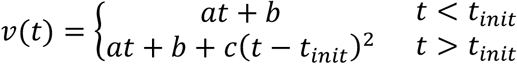

Here, *t*_*init*_ represents the time of response initiation. This function was fit to the region of the velocity profile spanning 100 ms before the target jump up to the point at which the velocity reached half of its peak value. To avoid spurious estimates of *t*_*init*_ under the Bimanual mapping, particularly during early blocks in which corrections were often weak, we constrained the parameter *c* to be greater than 10, to match the lowest estimates identified for the Baseline mapping. We fitted this model to averaged velocity profiles for each participant for a given target jump size and chunk of blocks after excluding extreme trials in which the cursor position at the time of the target jump deviated from the straight line to the target by more than 5 cm, or in which the absolute velocity of the cursor parallel to the target jump at the time of the target jump exceeded 0.2 m/s.

We determined the initial direction of movement of the cursor based on the direction of the smoothed velocity vector 100 ms after the detection of movement onset (based on a velocity threshold of 0.025 m/s). We determined the initial direction of movement of the left and right hand based on smoothed velocity vectors for each hand, calculated at the same time as cursor direction was determined. For trials in which either hand was stationary at that time (i.e., tangential velocity < 0.025m/s), initial direction was undefined for that trial and the trial was not included in relevant analyses.

### Statistics

Unless otherwise specified, we used two-tailed, paired t-tests to test for differences in behavior across time points and across conditions. For the control experiment, we assessed whether movement corrections depended on the control interface (left hand, right hand or both hands) using a one-way repeated measures ANOVA. To assess whether possible differences between one-hand and both-hands could account for the observed differences in correction speed between the Baseline and Bimanual mappings in Experiments 1 and 2, we compared the difference in behavior (the peak correction velocity and latency to peak correction velocity) between one-hand and both-hands condition in the control experiment and the Baseline and Bimanual mappings in Experiment 2 using a two-way ANOVA with experiment (Experiment 2 vs Experiment 3) and target-jump size (1.5 cm vs 3 cm) as factors. All statistical tests were conducted in MATLAB R2021a.

