## Supplementary Figures for "*De novo* motor learning of a bimanual control task over multiple days of practice"

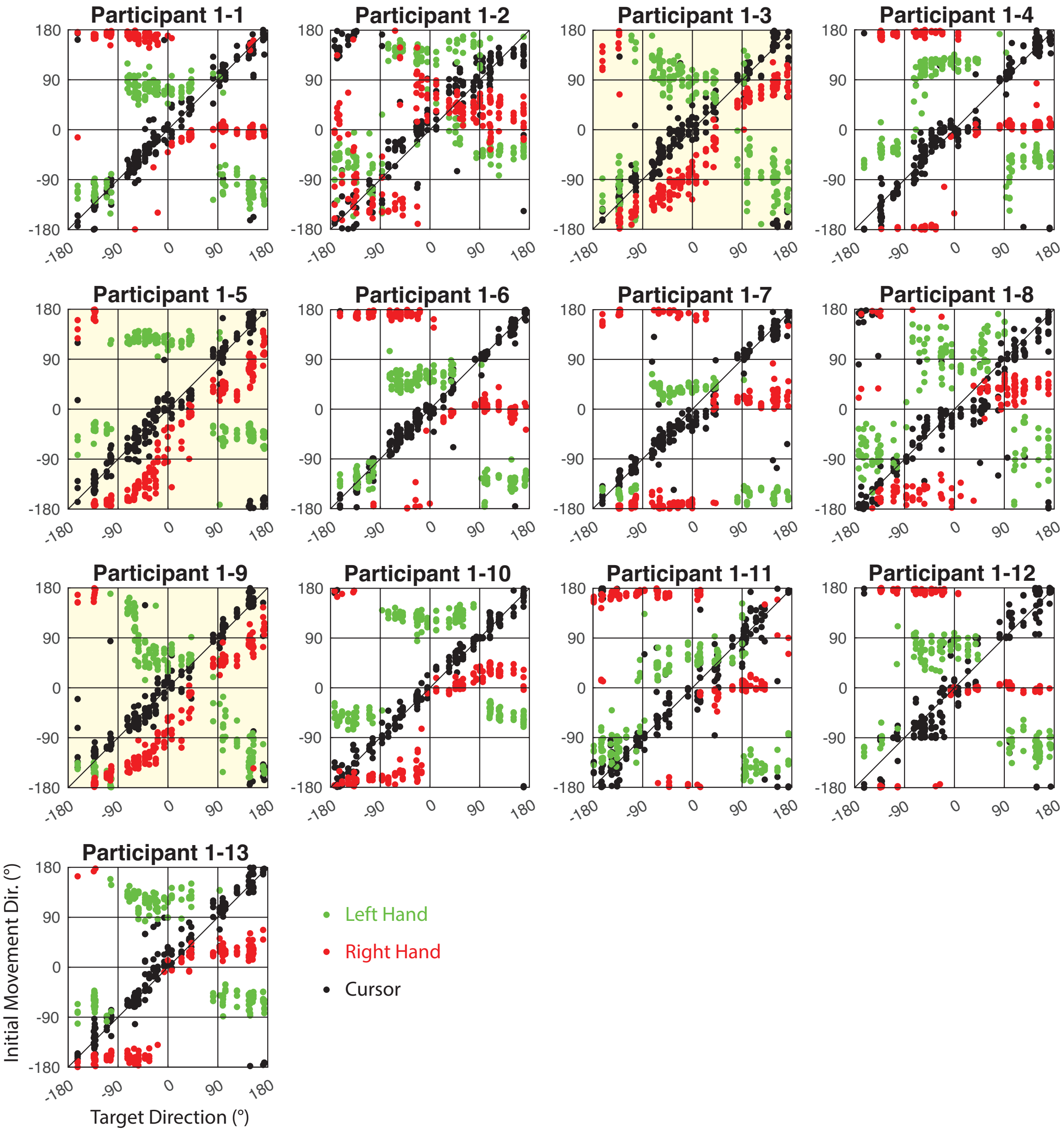


**Figure S1**. Learned coordination strategy for each participant in Experiment 1 (last 5 blocks). Green dots indicate initial direction of movement of the left hand in each trial as a function of initial direction of the target. Red dots indicate the same but for the right hand. Black dots indicate initial direction of the cursor. For trials in which only one hand moved initially, only one hand and the cursor are plotted. Three participants who exhibited an alternative strategy in which one hand was rotated 90° relative to the target are highlighted in yellow.

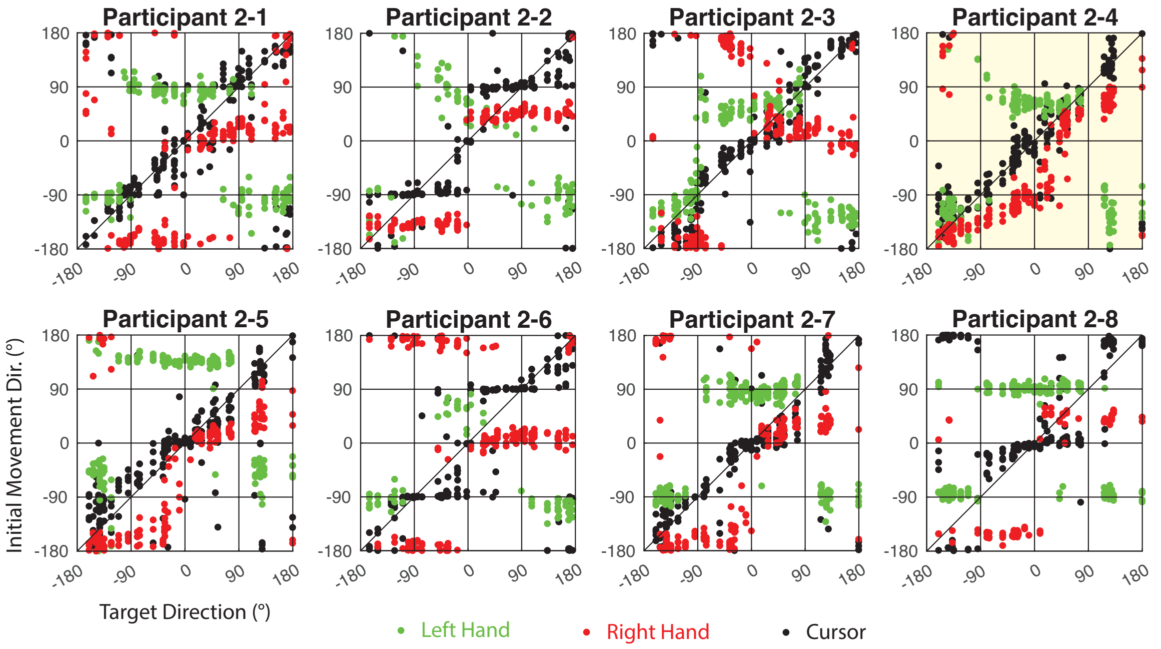


**Figure S2**. Learned coordination strategy for each participant in Experiment 2 (last 5 blocks). As Figure S1, except note that participants in Experiment 2 trained for only 3 days, rather than 4 days. Green dots indicate initial direction of movement of the left hand in each trial as a function of initial direction of the target. Red dots indicate the same but for the right hand. Black dots indicate initial direction of the cursor. For trials in which only one hand moved initially, only one hand and the cursor are plotted. One participant who exhibited an alternative strategy in which one hand was rotated 90° relative to the target is highlighted in yellow.
